# Genetic dissection of cell wall defects and the strigolactone pathway in Arabidopsis

**DOI:** 10.1101/586388

**Authors:** Vicente Ramírez, Markus Pauly

## Abstract

Defects in the biosynthesis and/or deposition of secondary plant cell wall polymers result in the collapse of xylem vessels causing a dwarfed plant stature and an altered plant architecture termed *irregular xylem (irx)* syndrome. For example, reduced xylan O-acetylation causes strong developmental defects and increased freezing tolerance. Recently, we demonstrated that the *irx* syndrome in the *trichome birefringence-like 29/eskimo1 (tbl29/esk1)* mutant is dependent on the biosynthesis of the phytohormone strigolactone (SL). In this report, we show that other xylan- and cellulose-deficient secondary wall mutants exhibit increased freezing tolerance correlated with the *irx* syndrome. In addition, blocking SL synthesis has also a suppressor effect on these phenotypes, suggesting a more general interaction between secondary wall defects and SL biosynthesis. In contrast, SLs do not play a role in developmental defects triggered by primary wall deficiencies, suggesting that the interaction is restricted to vascular tissue.

Through a reverse genetics approach the requirement of different components of the SL pathway impacting the *irx* syndrome in *tbl29* was evaluated. Our results are consistent with a specific role for carlactone in this process, and suggest that a MORE AXILLARY GROWTH 2 (MAX2)-independent SL perception mechanism might be involved.

## INTRODUCTION

Xylem elements are surrounded by a secondary wall composed mainly of cellulose, xylan and lignin, conferring unique physiochemical properties to these specialized cells. The functionality of the vascular system relies on the correct composition and structure of these modified walls, and most mutants affected in the biosynthesis and/or deposition of these three polymers show a collapsed vascular system (Turner and Somerville, 1997; Jones et al., 2001; Brown et al., 2005). These so-called *irregular xylem (irx)* mutants also exhibit severe growth defects including a dwarfed stature, reduced seed production, and an altered plant architecture.

In addition, several studies show that these developmental problems -globally termed as *irregular xylem* syndrome – are often accompanied by constitutive activation of stress responses (Xin and Browse, 1998; Chen et al., 2005; Keppler and Showalter, 2010). For example, reduction of xylan O-acetylation in *tbl29/esk1* mutant plants causes drastic xylem collapse accompanied by dwarfism and constitutive tolerance to freezing, drought and salt stresses (Xin and Browse, 1998; Xin et al., 2007; Lefebvre et al., 2011; Bouchabke-Coussa et al., 2008; Xiong et al., 2013). These phenotypes seem not to be a direct consequence of the defective secondary wall composition/structure, as two suppressor lines have been identified exhibiting rescued developmental- and stress-related phenotypes while the low xylan O-acetylation content was unchanged. In one case, *kaktus (kak)* mutant alleles suppress the *tbl29 irx* syndrome by boosting the vascular development resulting in normal xylem morphology and restoration of the water transport capacity (Bensussan et al., 2015). How *KAK*, a putative ubiquitin E3 ligase involved in endoreduplication and control of DNA content in trichomes, regulates xylem development remains elusive (Downes et al., 2003; El Refy et al., 2004; Bensusan et al., 2015). In another case, mutations in *MAX4* similarly rescues *tbl29* collapsed xylem, dwarfism and increased freezing tolerance (Ramirez et al., 2018). *MAX4* encodes a key enzyme in SL biosynthesis (Sorefan et al., 2003), suggesting that the tbl29-triggered *irx* syndrome is SL-dependent. This requirement was further confirmed by chemical complementation assays using a synthetic SL (i.e. GR-24). *tbl29 max4* double mutant plants growing in the presence of GR-24 exhibit collapsed xylem and dwarfed stature, indicating that complementing the SL-deficiency with exogenous applications of SL circumvent the *max4* effect (Ramirez et al., 20180).

SLs are a group of structurally similar compounds derived from carotenoids. Diverse SL compounds have been involved in the regulation of multiple processes in plants such as the interaction with root-parasitic plants and symbiotic arbuscular mycorrhizal fungi; developmental processes, including photomorphogenesis, root architecture, senescence, flower development or secondary growth; and adaptation responses to various biotic and abiotic stresses. However, the most extensively studied is the regulation of shoot branching (reviewed in Waters et al., 2017). The characterization of mutants showing a highly branched phenotype in Arabidopsis *(more axillary growth*; *max)*, pea (*ramosus*; *rms)*, rice *(high-tillering dwarf/dwarf; htd/d)* and petunia *(decreased apical dominance; dad)* has been responsible for the identification of multiple components involved in the SL biosynthesis and signal transduction pathway (Sorefan et al., 2003; Snowden et al., 2005; Arite et al., 2007; Booker et al., 2004; Johnson et al., 2006; Zou et al., 2006). In the first step of the SL biosynthesis, a plastid-localized β-carotene isomerase (AtD27 in Arabidopsis) converts all-trans-β-carotene into 9-cis-β-carotene, which is then used as substrate to generate carlactone by the consecutive action of two carotenoid-cleavage dioxygenase (CCD) enzymes, CCD7/MAX3/RMS5/D17/DAD3 and CCD8/MAX4/RMS1/D10/DAD1 (Schwartz et al., 2004; Alder et al., 2012; Waters et al., 2012). Carlactone seems to be the first active SL compound in the pathway and is thought to act as the precursor for other SL compounds (Scaffidi et al., 2013; Abe et al., 2014; Seto et al., 2014). The production of carlactone and its following conversion to carlactonic acid (in Arabidopsis likely performed by the MAX1 cytochrome P450 enzyme) seems to be a common strategy in plants. In contrast, it seems that the pathway diverges from there, and species-specific reactions transform carlactone into the more than 20 canonical and non-canonical SL compounds identified in plant exudates (Al-Babili and Bouwmeester, 2015; Tokunaga et al., 2015; Iseki et al., 2018). In Arabidopsis, carlactonic acid is transformed into methyl carlactonoate (MeCLA) by an unknown enzyme (Abe et al., 2014). MeCLA then binds and induces conformational changes to the D14 α/β- fold hydrolase allowing the interaction with a SKP1-CUL1-F-box-protein (SCF)-type ubiquitin ligase complex to transmit the SL signal via proteosomal degradation of negative regulators of SL signaling. An integral component of this complex, the MAX2 leucine rich F-box protein, seems to be involved in the ubiquitination of putative transcriptional regulators controlling most of the SL-dependent responses characterized so far, including members of the SUPPRESSOR OF MAX2 (SMAX1) and SMAX1-LIKE families, the BRI1-EMS SUPPRESSOR1 (BES1) or the DELLA family (Nakamura et al., 2013; Wang et al., 2013; Jiang et al., 2013; Soundappan et al., 2015; Wang et al., 2015). However, the function and exact mechanism of action of most of these target proteins is still under debate. Upon D14-MeCLA interaction, D14 returns to the initial conformation inducing the hydrolytic degradation of MeCLA and restoring the system after signal transmission (Seto et al., 2019).

## RESULTS

### Several cellulose- and xylan-deficient mutants show increased freezing tolerance

Similar to reduced xylan O-acetylation in the *tbl29* Arabidopsis mutant, other secondary cell wall defects impact xylem morphology, plant size, and architecture. Because the *tbl29* mutant exhibits freezing tolerance, a potential general correlation of the irregular xylem phenotype with increased freezing tolerance was investigated (Figure 1). For this purpose, Arabidopsis T-DNA insertion lines were obtained for genes involved in the synthesis of cellulose, xylan and lignin. The *irregular xylem 1 (irx1)* and *irregular xylem 3 (irx3)* mutants have a severe deficiency in the deposition of cellulose in secondary walls, caused by mutations in the *CELLULOSE SYNTHASE 8* and *7* genes, respectively (Taylor et al., 2000). The *irregular xylem 9 (irx9)* and *parvus* mutants are affected in the synthesis of the β-1-4-xylan backbone and the tetrasaccharide structure located at the reducing end of xylan, respectively (Bauer et al., 2006; Brown et al., 2007; Lee et al., 2007a; Lee et al., 2007b; Peña et al., 2007). Mutant alleles of *CINNAMOYL COA REDUCTASE 1 (IRX4/CCR1)* and *CAFFEOYL SHIKIMATE ESTERASE (MAGL3/CSE/LYSOPL2)* genes affect lignin deposition (Jones et al., 2001; Vanholme et al., 2013). All of these mutants share with *tbl29* various developmental defects including reduced rosette size, a dwarfed stature and a characteristic dark green color in the leaves (Figure 1). When freezing tolerance was evaluated, all tested alleles of cellulose-*(irx1* and *irx3*, Figure 1A-B) and xylan-compromised mutants *(irx9* and *parvus;* Figure 1C-D) showed an increase in freezing tolerance compared to their respective controls. Although the *tbl29* mutants showed the highest survival rate (>90%) under the freezing conditions used, *irx1, irx3, irx9*, and *parvus* mutant lines showed also high values (>70%) compared to the respective controls (<10% survival rate). Interestingly, the two lignin-compromised mutants analyzed (i.e. *irx4* and *magl3/cse;* Figure 1E-F) showed a wildtype-like response in our freezing assay despite dwarfism and collapsed vessel elements (Jones et al., 2001; Vanholme et al., 2003). These results could be reproduced in a second Arabidopsis accession (Landsberg (Laer)), at least for *irx3, irx4* and *parvus*, where mutant alleles were available.

**Figure 1.**
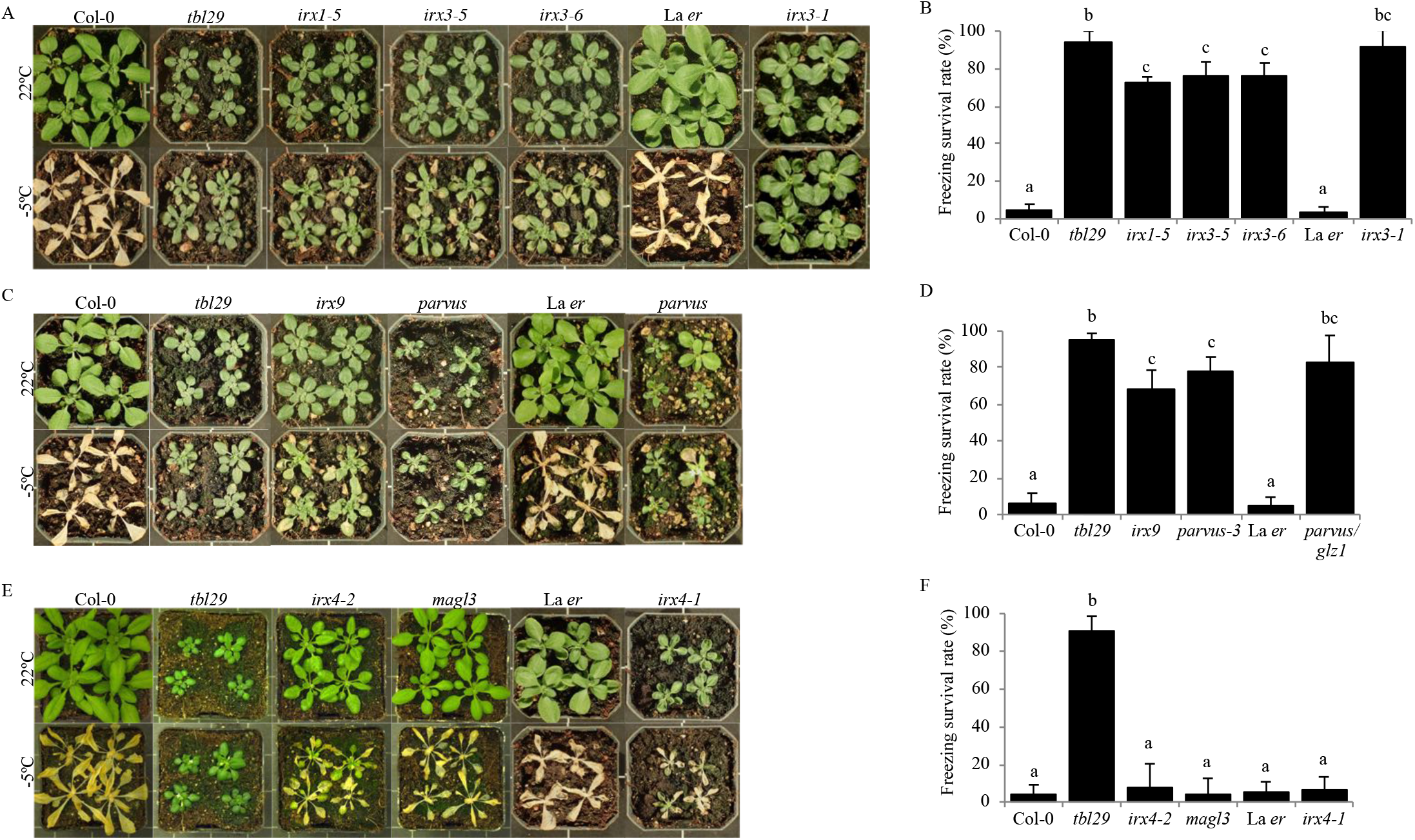
Freezing tolerance in irregular xylem mutants. (A, C and E) Representative pictures of 4-week-old plants of the indicated genotypes before (upper panel) and after (lower panel) freezing treatment. (B, D and F) Survival rate after freezing assay. Data are represented as mean (AVG) ± the standard deviation (SD) of 3 independent experiments (≥20 plants/experiment). Means with different letters are significantly different (Tukey’s HSD, p<0.05) in the significance (S) column.

### Effect of SL deficiency on other secondary wall mutants

Mutations in the *MAX4* SL-biosynthetic gene rescues the dwarfism, collapsed xylem and increased freezing tolerance phenotypes associated with defects in xylan-*O*-acetylation in *tbl29* (Ramirez et al., 2018). The effect of the *max4* mutation on the phenotype of freezing tolerant cellulose-(i.e. *irx1* and *irx3)* and xylan-(i.e. *irx9* and *parvus)* deficient mutants was analyzed (Figure 2 and Figure 3). Detailed analyses of the growth habit of the respective double mutants generated showed that the presence of *max4* leads to an increase in plant size in *irx1 max4*, *irx3 max4, irx9 max4* and *parvus max4* plants. However, these double mutants do not exhibit a fully restored wildtype growth as was the case in *tbl29 max4* plants (Figure 2A-B and Figure 3A-B). The increase in plant size is also accompanied by an increase in the number of rosette branches in the mutants (Figure 2C and Figure 3C). A modest improvement is also observed in the collapsed xylem morphology in stem sections (Figure 2D and Figure 3D). Although most of the xylem still remains irregular, more larger vessels are observed in the double mutants compared to the single mutants.

**Figure 2.**
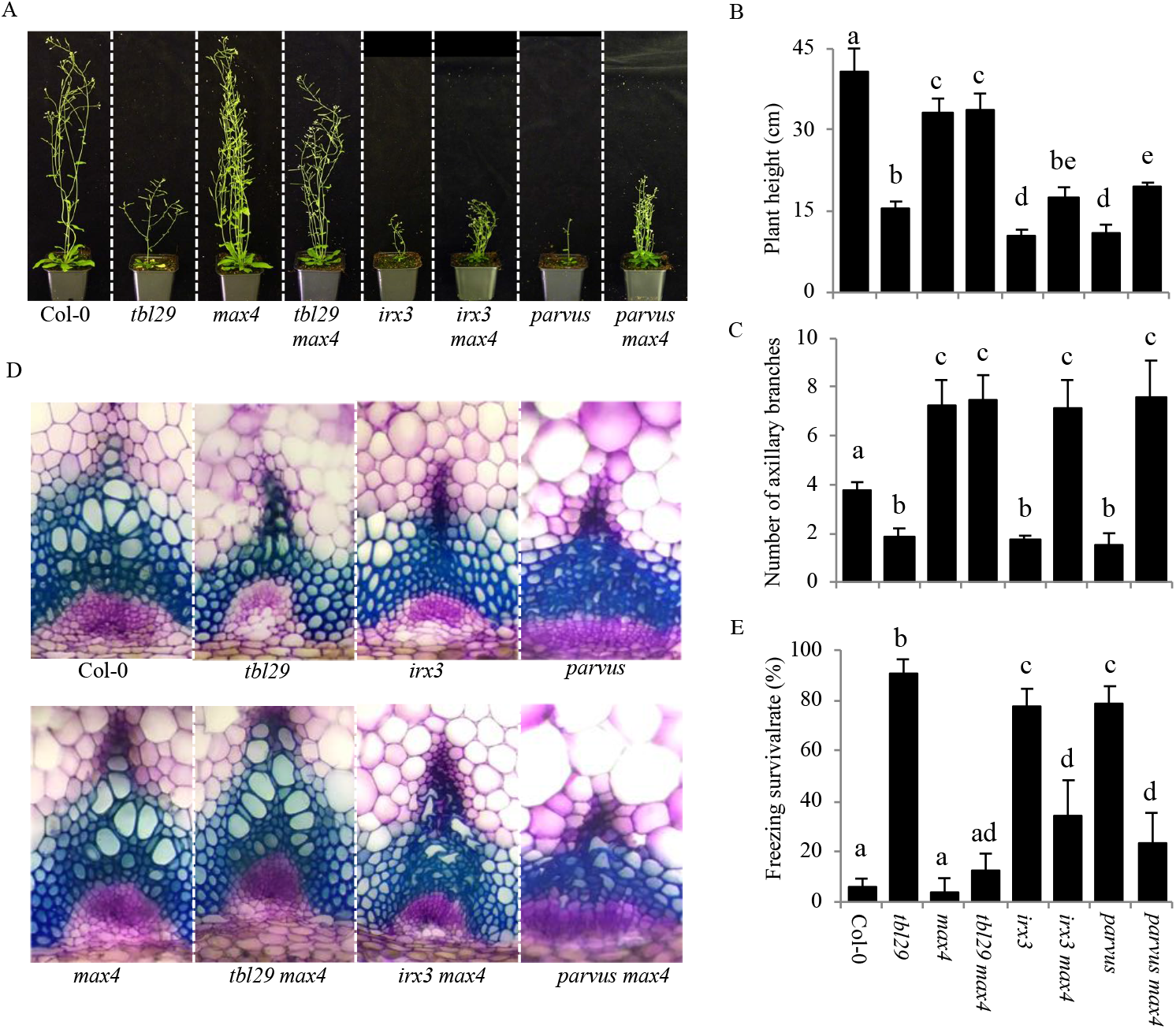
*max4* mutation in other secondary cell wall mutants. (A) Growth phenotypes of 6-week-old plants. (B) Primary inflorescence heights (cm) (C) Number of rosette branches (D) Toluidine-O-Blue stained cross section of inflorescence stems. (E) Survival rate of freezing assay. Means with different letters are significantly different (Tukey’s HSD, p<0.05).

Similarly, the freezing tolerance in *irx1 max4, irx3 max4* and *parvus max4* is significantly reduced compared to *irx1, irx3* and *parvus* although the recovery is not complete as indicated by the intermediate survival rates of the double mutants (Figure 2E and Figure 3E). In the case of *irx9* and *irx9 max4* no significant reduction in freezing tolerance was observed.

**Figure 3.**
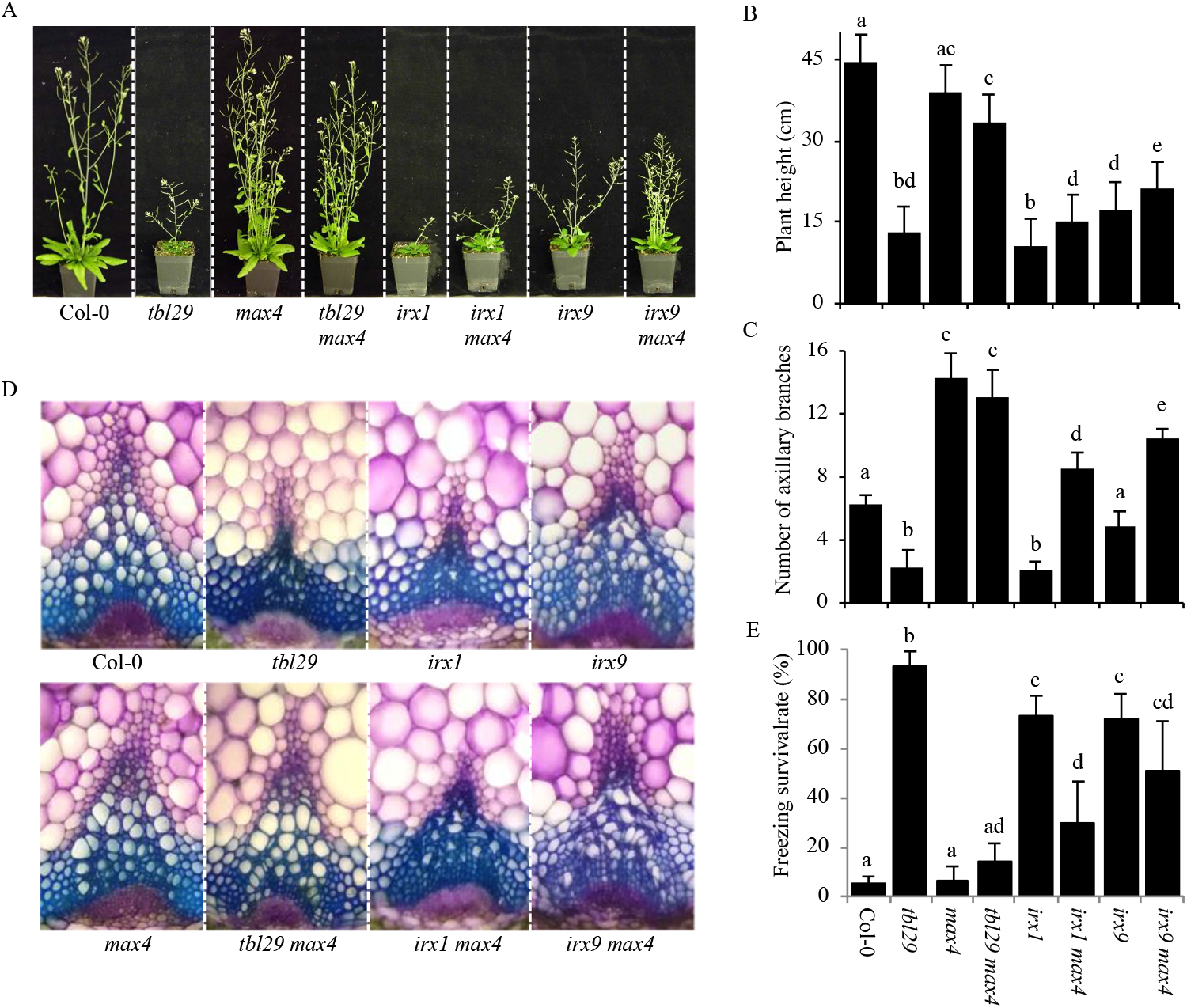
*max4* mutation in other secondary cell wall mutants. (A) Growth phenotypes of 6-week-old plants. (B) Primary inflorescence heights (cm) (C) Number of rosette branches (D) Toluidine-O-Blue stained cross section of inflorescence stems. (E) Survival rate of freezing assay. Means with different letters are significantly different (Tukey’s HSD, p<0.05).

Wall analyses showed no obvious differences in the composition of matrix polysaccharides caused by the introduction of *max4* into the various mutant backgrounds (Supp. Tables 1 and 2), and the defects caused by the *irx* mutations remain intact in the respective double mutants. However, a minor increase in cellulose content was noticed in *tbl29 max4, irx1 max4, irx3 max4, irx9 max4* and *parvus max4* compared to the respective *tbl29, irx1, irx3, irx9* and *parvus* single mutants.

Taken together, these results indicate that although *max4* is able to override the consequences of xylan hypoacetylation on plant development and freezing tolerance, it is only able to partially compensate for the more severe wall defects in the xylem vessels of reduced cellulose and altered xylan mutants.

### Effect of SL deficiency on primary wall mutants

Defects in primary wall cellulose biosynthesis also lead to developmental defects including dwarfism of etiolated hypocotyls (Fagard et al., 2000; Scheible et al., 2001). In order to evaluate the effect of blocking SL synthesis on primary wall-deficient mutants, the *max4* mutation was introduced in two mutant alleles of the *PROCUSTE 1/ISOXABEN RESISTANT 2/CELLULOSE SYNTHASE 6* gene *(PRC1/IXR2/CesA6).* The *prc1-1* strong allele shows severe reductions in cellulose deposition in the primary wall. As a result, seedling length is reduced by more than 80% compared to the wildtype. A weaker allele, *ixr2-1*, only shows a mild reduction of seedling growth (≈25%). Introduction of *max4* mutation in *prc1-1* or *ixr2-1* mutant backgrounds does not complement the developmental defects, as the seedling length of double mutant plants remains reduced (Figure 4A-B). As expected, the presence of *max4* does not change the crystalline cellulose content on any of the two mutant backgrounds (Figure 4C). These results suggest that the dwarfism triggered by primary wall deficiencies is SL-independent.

**Figure 4.**
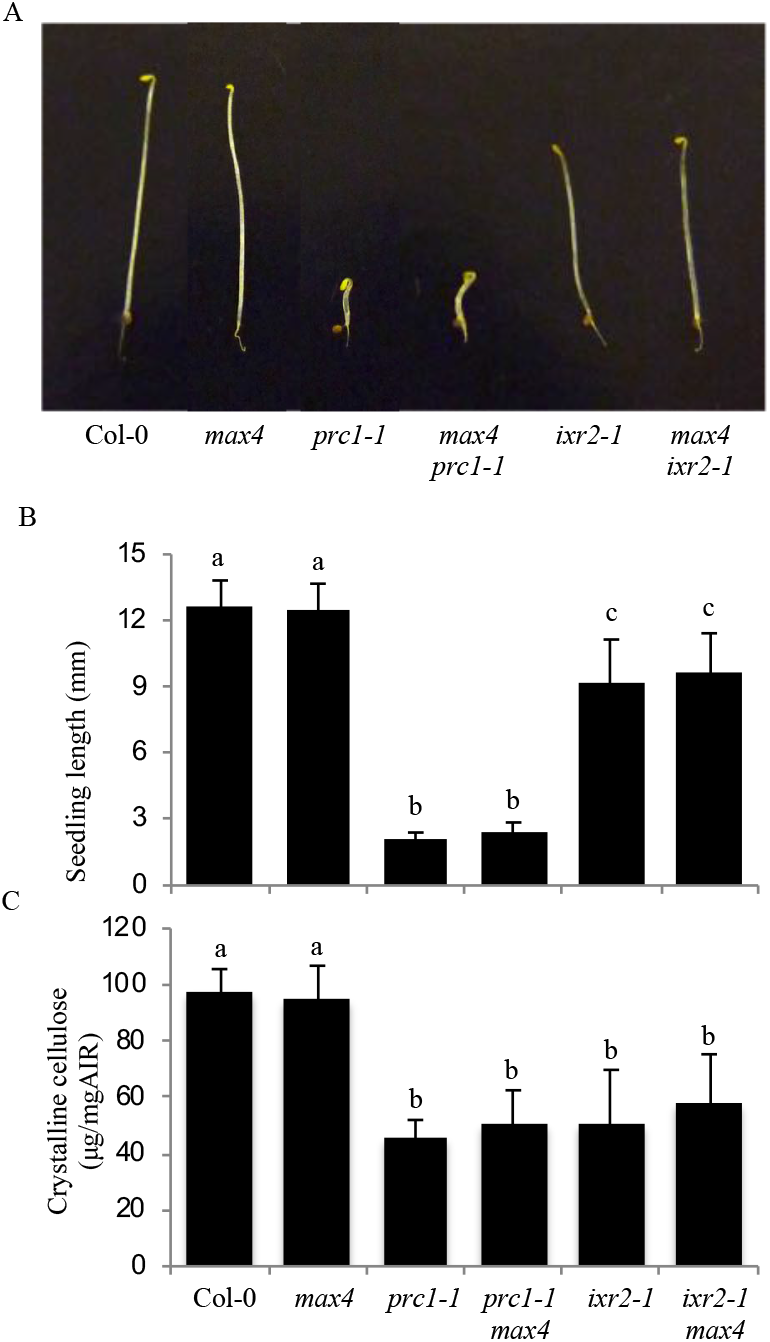
*max4* mutation in primary wall-deficient mutants. (A) Representative pictures of dark-grown 5-day-old seedlings. (B) Seedling length. Bars represent mean (AVG) ± the standard deviation (SD) of individual seedlings (n≥100). (C) Crystalline cellulose content. Bars represent mean (AVG) ± the standard deviation (SD) of n=5 biological replicates. Means with different letters are significantly different (Tukey’s HSD, p<0.05)

### Effect of SL biosynthesis- and perception-deficient mutants on *tbl29*-associated irregular xylem phenotype

The effect of blocking different steps of the SL pathway was investigated by introducing mutant alleles of *MAX1, MAX2*, and *MAX3* genes into the *tbl29* mutant background to generate the corresponding *tbl29 max1, tbl29 max2* and *tbl29 max3* double mutants. A detailed characterization of these plants including the previously characterized *tbl29 max4* double mutant line was performed in terms of plant growth habit, xylem morphology and freezing tolerance (Figure 5). Plant height in *tbl29 max3*, and *tbl29 max4* double mutant was comparable to the *max3* and *max4* single mutants, rescuing almost completely the *tbl29* dwarf phenotype. On the other hand, *tbl29 max1* and *tbl29 max2* double mutant plants were still dwarfed, with stem heights similar to *tbl29* plants. This result was confirmed with two different *max2* mutant alleles (i.e. *max2-1* and *max2-2).*

**Figure 5.**
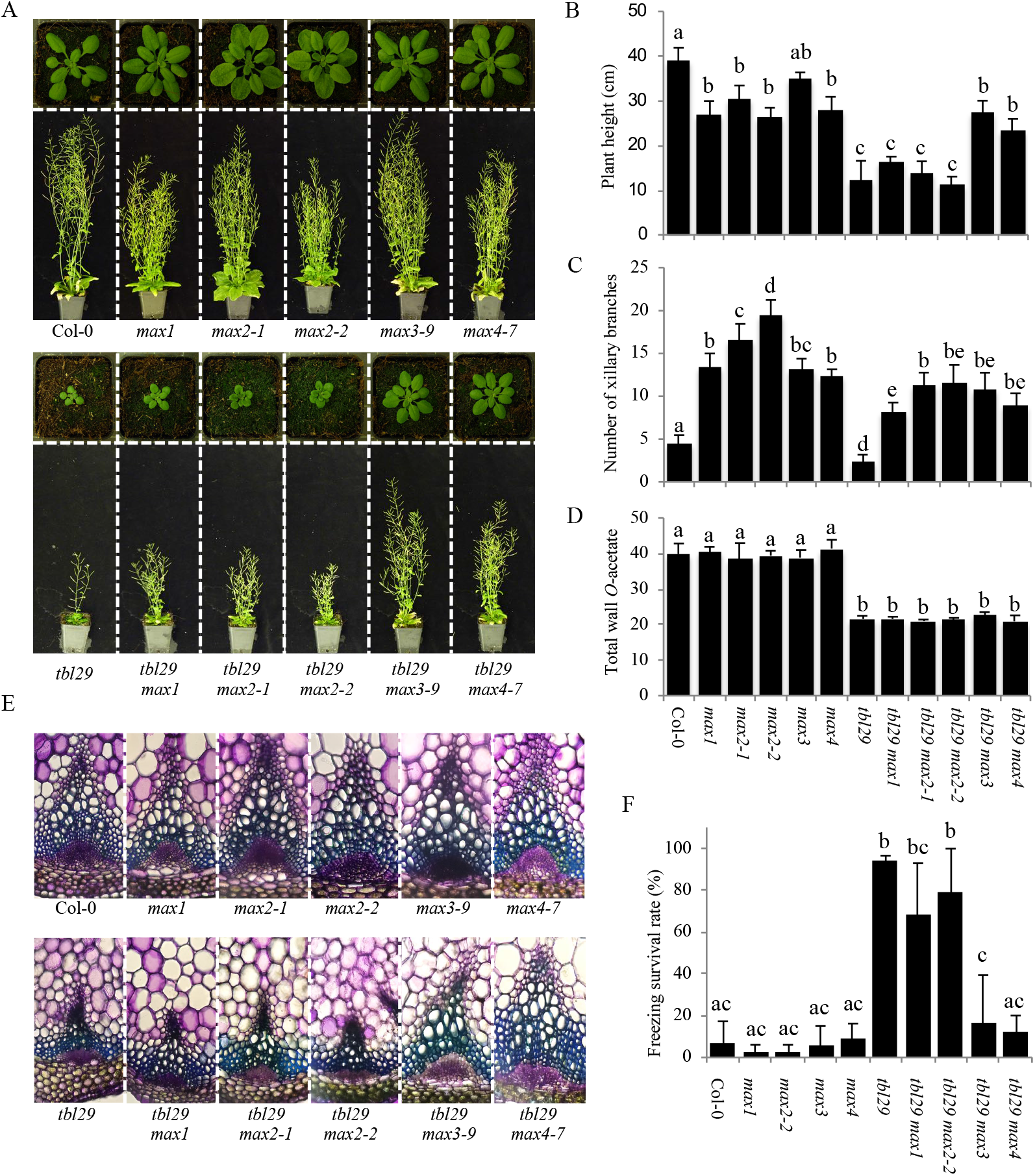
*max* mutations in tbl29. (A) Growth phenotypes of 4-(upper panel) and 6-week-old (lower panel) plants. (B) Primary inflorescence heights (cm) (C) Number of rosette branches (D) Toluidine-O-Blue stained cross section of inflorescence stems. (E) Survival rate of freezing assay. Means with different letters are significantly different (Tukey’s HSD, p<0.05).

Toluidine blue staining of stem sections of *max1, max2, max3* and *max4* single mutants showed no obvious differences with Col-0. While *tbl29 max3* and *tbl29 max4* double mutant plants also showed normal xylem development, equivalent stem sections of *tbl29 max1, tbl29 max2-1* and *tbl29 max2-2* double mutant plants showed a collapsed xylem comparable to *tbl29* single mutant (Figure 5E). As previously shown in for *tbl29 max4*, the rescued developmental defects in *tbl29 max3* are not associated with the recovery of normal xylan O-acetylation levels as both double mutants show reduced values comparable to *tbl29* single mutant (Figure 5D).

No significant differences were found in the response to freezing temperatures of *max1, max2, max3* and *max4* single mutants compared to Col-0, with survival rates below 15%. In contrast, *tbl29 max1* and *tbl29 max2* plants were highly tolerant to freezing temperatures, with a survival rate over 70% similarly to *tbl29*. As with plant height, the increased freezing tolerance associated to *tbl29* was suppressed in *tbl29 max3* and *tbl29 max4* double mutants with a survival rate comparable to Col-0 (Figure 5F).

As previously reported *tbl29* mutant plants exhibited a reduced number of axillary branches compared to Col-0. On the other hand, *max* mutants showed an increased branching phenotype due to SL-deficiency (Sorefan et al., 2003). All *tbl29 max* double mutants generated showed significantly increased number of axillary branches compared *tbl29*, regardless of whether the plant height was rescued or not (Figure 5C). This result strongly suggests that the suppression of *tbl29* phenotypes observed in *tbl29 max3* and *tbl29 max4* is not due to an indirect effect of a change in plant architecture (i.e. increased number of axillary branches). Together, these results indicate that blocking any of the two carotenoid cleavage dioxygenases (i.e. MAX3 and MAX4) involved in carlactone biosynthesis rescues the collapsed xylem, growth defects and increased freezing tolerance associated to the reduction of xylan *O*-acetylation caused by *TBL29* loss of function. On the other hand, mutations in genes encoding downstream enzymes in the SL pathway failed to complement the *tbl29* mutant phenotypes, suggesting that production (i.e. MAX1) or perception (i.e. MAX2) of other SL compounds are not required.

## DISCUSSION

### Freezing tolerance in secondary wall-deficient mutants

A side-by-side analysis of multiple secondary wall-deficient mutants showed that increased freezing tolerance seems to be a common phenotype of plants with altered xylan composition and cellulose content (Figure 1). The increased freezing tolerance of *tbl29/esk1* mutant alleles has been known for 20 years (Xin and Browse, 1998). Since then, various *tbl29* stress-related phenotypes have been the focus of several studies (Xin et al., 2007; Lefebvre et al., 2011; Bouchabke-Coussa et al., 2008). Similarly, various reports have also shown stress-related phenotypes for individual secondary wall-deficient mutants such as increased tolerance to salt stress, drought or pathogen attack (Chen et al., 2005; Keppler and Showalter, 2010; Hernández-Blanco et al., 2007; Ramirez et al., 2011). Although the mechanism by which *tbl29/esk1* plants become freezing tolerant is unknown, evidence has been presented that it is independent from the classical ABA hormonal pathway or the ICE1–CBF cold response pathway (Lefebvre et al., 2011). The current hypothesis entails that the increased freezing tolerance (and other constitutive stress responses) in *tbl29/esk1* mutants are caused by a transpiration imbalance produced by the collapse of the xylem. The constitutive freezing tolerance phenotype of multiple secondary wall mutants suggest that the same mechanism could apply to xylan- and cellulose-deficient mutants. Interestingly, the two lignin-deficient mutants analyzed here exhibited a wildtype-like response in our freezing test, suggesting that lignin defects leading to xylem collapse might not trigger the same mechanism.

### Effect of SL-deficiency in primary and secondary wall mutants

Recently, we showed how mutations in the *MAX4* SL-biosynthetic gene where able to suppress the developmental- and stress-related phenotypes caused by the reduction in the xylan *O*-acetylation in *tbl29* (Ramirez et al., 2018). The severity of the *irx* syndrome in *tbl29* is similar to other xylan- and cellulose-deficient mutants as *irx1, irx3*, or *parvus* in terms of dwarf stature, collapsed xylem or increased freezing tolerance (Figure 2 and Figure 3). Introducing the *max4* mutation in these secondary wall mutants has a positive effect in plant height and collapsed xylem and partially rescues the constitutive freezing tolerance compared to the single mutants (Figure 2 and Figure 3). These data demonstrate that, in general, secondary wall polysaccharide defects trigger the SL-pathway without repairing the defective wall structures or other structural compensations. However, although blocking SL biosynthesis recovers the *irx* syndrome caused by a reduction in xylan *O*-acetylation, it is not able to restore completely the more severe secondary wall defects such as reduced cellulose content or defects in xylan backbone biosynthesis. It thus seems that in the case of minor structural modifications (i.e. reduced xylan *O*-acetylation), blocking SL biosynthesis is able to short circuit the wall defect-triggered signal and rescue the irregular xylem-derived phenotypes. In contrast, when the defect compromises the secondary wall integrity/functionality (i.e. defects in the xylan backbone or cellulose synthesis) the rescue is only partial.

Multiple lines of evidence indicate the existence of a mechanism monitoring the integrity of primary walls (reviewed by Wolf, 2017). Primary wall-deficient mutants display retarded growth due to defective cellulose deposition and can be complemented by blocking components of a still not completely understood signaling network without affecting the wall composition (Fagard et al., 2000; Scheible et al., 2001; Wolf, 2017). Similar to the case of *tbl29*, it seems that the developmental phenotypes exhibited by these primary wall mutants (e.g. *prc1* or *ixr2*) are not a direct consequence of the wall defect but the activation of a downstream signaling response. Introduction of *max4* mutation in strong and mild mutant alleles of a primary wall-specific cellulose synthase (i.e. *CesA6)* does not complement the developmental defects (Figure 4), suggesting that in the primary cell wall integrity system the SL pathway is not involved, but the SL pathway has a secondary wall-specific function likely restricted to vascular tissue.

### Specificity of *max* mutations as suppressors of *tbl29*

While *max3* and *max4* mutations were able to suppress dwarfism, collapsed xylem, and increased freezing tolerance associated to *tbl29*, other mutations causing SL deficiency *(max1)* or SL insensitivity *(max2)* failed to complement (Figure 5). Considering that all *max* mutations induced an increased axillary shoot branching in the *tbl29* mutant background regardless of the suppression of the *tbl29*-associated phenotypes, we can conclude that this phenomenon is not due to an altered plant architecture induced by a SL deficiency. Our genetic data suggests that specifically some components of the SL biosynthetic pathway are involved. MAX3 and MAX4 encode two carotenoid-cleavage dioxygenase enzymes, whose sequential actions produce carlactone from β-carotene (Matusova et al., 2005; Auldridge et al., 2006; Alder et al., 2012). According to our results, endogenous production of carlactone would be required for the expression of developmental-and stress-related phenotypes caused by certain secondary wall defects (e.g. *tbl29).* Intriguingly, it seems that this effect is independent on downstream steps in the SL biosynthesis such as MAX1 and MAX2. These results apparently contradict the current SL model of action, based on the SL-dependent inhibition of shoot branching. Our genetic data imply that carlactone has a regulatory role independent on the perception of SL compounds through MAX2. However, this is not the first result contradicting this model. For example, carlactone applications are not only able to complement the lateral inflorescence phenotype in SL-deficient mutants (e.g. *max1* and *max4)* but also partially in *max2* SL-insensitive mutant plants, suggesting the existence of a MAX2-independent perception mechanism for carlactone. In contrast, MeCLA and CLA applications complement only SL-deficient mutants but not in *max2*, indicating that unlike carlactone, MeCLA/CLA perception requires MAX2 (Abe et al., 2014). Carlactone has been proposed as a precursor of other SL compounds, and subsequent species-specific rearrangements and modifications catalyzed by downstream enzymes would convert carlactone into the large diversity of SL compounds found in plants (Seto et al., 2014; Iseki et al., 2018). Thus, we cannot rule out the possibility that not carlactone itself, but a carlactone-derived, MAX1-independent SL compound is involved in this regulation of xylem development in secondary wall deficient mutants.

## MATERIALS AND METHODS

### Growth conditions

Seeds were stratified in 0.2% Agarose for 4 days in the dark at 4°C. After stratification, seeds were sown on soil and grown in environmentally controlled growth chambers (8,000 luxes, 16 hr light, 22°C/8 hr dark, 19°C).

### Mutant genotyping

Double mutants were generated by crossing and identified by genotyping of individuals in the F2 generation. Genotyping of T-DNA lines was performed as described in Ramirez et al., 2018 (see Supp. Table 3 for primer sequences). To genotype *max2-1* mutation, the amplicon generated by PCR using *max2-1 F/ max2-1 R* primer combination was digested with *ApoI* restriction enzyme resulting in a 147 bp single fragment in wildtype plants and two fragments (76 and71 bp) in mutant plants. *max3-9* genotyping was performed using *max3-9 F/max3-9 R* primer combination, resulting in 120 bp PCR amplicon in wildtype plants and 131 bp in the mutant. To genotype the EMS-derived *max1* and *max2-2* mutations, PCR reactions with *max1 F/max1 R* and *max2-2 F/max2-2 R* primer combinations were performed and the respective amplicons sequenced. Genotyping of prc1-1 was done with *pcr1-1 F/pcr1-1 R* primer combination and further digestion with *HpyCH4V* restriction enzyme resulting in 3 fragments in the wildtype (135, 70 and 56 bp) and only two in the mutant (135 and 126 bp). ixr2-1 genotyping was done using the following primer combinations: *ixr2-1* Fin/ *ixr2-1* Rout, resulting in amplification only in wildtype plants; *ixr2-1* Fout/ *ixr2-1* Rin, resulting in amplification only in mutant plants; and *ixr2-1* Fout/ *ixr2-1* Rout, resulting in amplification in all individuals. Primer sequences are described in Supp. Table 4.

The different mutant lines used in this study were obtained from the NASC (http://arabidopsis.info) (Scholl et al., 2000) and ABRC (https://abrc.osu.edu) Arabidopsis stock centers. Stock numbers for each mutant line are described in Supp. Table 5.

### Freezing experiments

4-week-old plants of the indicated genotypes were incubated at −5°C in a Coolfreeze CF-110 cooling box (Waeco) for 20 hr. After the freezing treatment, plants were transferred to the growth chamber. Survival rates were calculated as the % of plants alive after 3 days. Individual pictures were taken before and after the freezing exposure using a Lumix DMC-FZ35 camera (Panasonic)

### Plant measurements

Plant height and the number of axillary branches were recorded on 6-week old plants. Individual pictures were taken using a Lumix DMC-FZ35 camera (Panasonic).

Seedling length measurements were performed as described in Fagard et al., 2000 with minor modifications. Briefly, seeds were surface-sterilized in a solution of 70% ethanol and 0.01% Tween-20 and sown in half strength Murashige and Skoog-containing media without sucrose (Duchefa). Plates were incubated for 2 days at 4°C, then exposed to light for 1 hour to synchronize germination, covered in two layers of aluminum foil and incubated at 24°C for 4 days. Hypocotyls were horizontally transferred to agar plates and their image was recorded with a Lumix DMC-FZ35 camera (Panasonic). Hypocotyl lengths were measured using ImageJ free software.

### Xylem morphology

Xylem morphology was evaluated as described in Ramirez et al., 2018. Equivalent segments of plant stems from the various genotypes were used to prepare hand-cut sections. After incubation in 0.02% Toluidine Blue O solution (Sigma-Aldrich) for 2 minutes, sections were washed three times with sterile water and observed under a bright-field lighting microscope (Leica DM2000 LED). At least 10 plants/genotype and 20 sections/plant were evaluated.

### Cell wall composition

Primary stems from individual 6-week-old plants were cut in 1cm segments and ground for 2 minutes in a MM400 mixer mill (Retsch Technology) after lyophilization using a ScanVac CoolSafe Freeze-dryer (Labogene). Total destarched alcohol insoluble residue was prepared and cellulose content, matrix polysaccharide composition and total wall acetate was determined according to Foster et al., 2010 as described in Ramirez et al., 2018.

## Author contributions

M.P. and V.R. designed the research; V.R. conducted experiments; M.P. and V.R. wrote the paper

## Supplemental Data files

Supplemental Table 1. Effect of *max4* mutation in the cell wall composition of *irx3* and *parvus*.

Supplemental Table 2. Effect of *max4* mutation in the cell wall composition of *irx1* and *irx9.*

