## Supplemental data for "Genetic dissection of cell wall defects and the strigolactone pathway in Arabidopsis"

| | Fuc | | | Rha | | | Ara | | | Gal | | | Glc | | | Xyl | | | Man | | | GalA | | | GlcA | | | Cellulose | | | Acetate | | |
|---|---|---|---|---|---|---|---|---|---|---|---|---|---|---|---|---|---|---|---|---|---|---|---|---|---|---|---|---|---|---|---|---|---|
| | AVG | SD | S | AVG | SD | S | AVG | SD | S | AVG | SD | S | AVG | SD | S | AVG | SD | S | AVG | SD | S | AVG | SD | S | AVG | SD | S | AVG | SD | S | AVG | SD | S |
| **Col-0** | 1.5 ± | 0.2 | a | 7.6 ± | 0.8 | a | 5.8 ± | 1.2 | a | 10.2 ± | 1.2 | a | 14.0 ± | 1.3 | a | 97.2 ± | 5.2 | a | 9.1 ± | 1.1 | a | 29.0 ± | 4.2 | a | 2.7 ± | 0.5 | ad | 361.1 ± | 12.4 | ab | 45.9 ± | 1.5 | a |
| ***tbl29*** | 1.5 ± | 0.1 | a | 9.8 ± | 0.7 | c | 9.1 ± | 1.1 | c | 12.0 ± | 1.0 | a | 16.6 ± | 1.8 | b | 102.9 ± | 11.7 | b | 11.1 ± | 1.0 | b | 30.1 ± | 1.2 | a | 3.6 ± | 0.7 | ac | 309.7 ± | 33.5 | b | 27.3 ± | 1.5 | bd |
| ***tbl29 max4*** | 1.3 ± | 0.5 | a | 7.0 ± | 1.7 | a | 5.7 ± | 1.4 | a | 9.0 ± | 2.2 | a | 15.1 ± | 3.5 | b | 90.7 ± | 14.0 | a | 8.6 ± | 1.4 | a | 29.7 ± | 1.0 | a | 4.7 ± | 0.8 | bc | 364.6 ± | 33.2 | a | 24.2 ± | 2.8 | b |
| ***irx3*** | 1.7 ± | 0.2 | a | 11.9 ± | 0.6 | b | 19.8 ± | 2.2 | d | 15.4 ± | 1.1 | b | 9.5 ± | 1.5 | ac | 159.5 ± | 10.6 | c | 6.4 ± | 0.4 | c | 38.8 ± | 2.5 | b | 4.3 ± | 1.1 | c | 91.6 ± | 15.5 | c | 72.6 ± | 3.7 | c |
| ***irx3 max4*** | 1.9 ± | 0.2 | a | 11.7 ± | 0.8 | b | 13.7 ± | 2.7 | b | 14.3 ± | 1.4 | b | 10.5 ± | 1.0 | c | 155.8 ± | 12.6 | c | 7.2 ± | 0.4 | c | 37.1 ± | 4.6 | b | 5.5 ± | 0.9 | c | 120.5 ± | 7.6 | d | 73.7 ± | 4.4 | c |
| ***parvus*** | 2.5 ± | 0.2 | b | 11.2 ± | 0.4 | bc | 13.1 ± | 0.9 | b | 21.1 ± | 1.4 | c | 21.1 ± | 4.2 | b | 47.7 ± | 2.7 | d | 16.6 ± | 1.1 | d | 35.3 ± | 2.6 | b | 1.0 ± | 0.4 | d | 309.4 ± | 14.9 | b | 31.7 ± | 1.5 | d |
| ***parvus max4*** | 2.6 ± | 0.1 | b | 11.6 ± | 0.3 | b | 13.7 ± | 1.1 | b | 20.7 ± | 1.2 | c | 20.2 ± | 3.7 | b | 46.2 ± | 3.4 | d | 15.7 ± | 1.6 | d | 35.5 ± | 1.7 | b | 1.4 ± | 0.4 | d | 327.7 ± | 9.1 | e | 31.6 ± | 1.9 | d |

**Supp. Table 1. *Effect of max4* mutation in the cell wall composition of *irx3* and *parvus*.** Monosaccharide composition, cellulose and acetate content of stem cell walls. Data are represented as mean (AVG) ± the standard deviation (SD) of biological replicates (n=5). Means with different letters are significantly different (Tukey's HSD, $p<0.05$) in the significance (S) column.

Fuc=Fucose; Rha=Rhamnose; Ara=Arabinose; Gal=Galactose; Glc=Glucose; Xyl=Xylose; Man=Mannose; GalA; Galacturonic Acid; GlcA= Glucuronic Acid

| | Fuc | | | Rha | | | Ara | | | Gal | | | Glc | | | Xyl | | |
|---|---|---|---|---|---|---|---|---|---|---|---|---|---|---|---|---|---|---|
| | AVG | ± SD | S | AVG | ± SD | S | AVG | ± SD | S | AVG | ± SD | S | AVG | ± SD | S | AVG | ± SD | S |
| **Col-0** | 1.3 | ± 0.1 | a | 8.9 | ± 0.2 | a | 7.6 | ± 0.5 | a | 13.7 | ± 0.6 | a | 17.8 | ± 1.6 | a | 92.3 | ± 9.9 | a |
| ***tbl29*** | 1.3 | ± 0.1 | a | 10.1 | ± 0.8 | b | 9.1 | ± 0.9 | a | 14.6 | ± 1.2 | a | 17.5 | ± 3.5 | a | 94.6 | ± 9.2 | a |
| ***max4*** | 1.3 | ± 0.1 | a | 8.7 | ± 0.4 | ac | 7.6 | ± 0.6 | a | 13.7 | ± 0.8 | a | 18.9 | ± 2.2 | a | 100.5 | ± 6.2 | a |
| ***tbl29 max4*** | 1.2 | ± 0.1 | a | 8.2 | ± 0.7 | ac | 6.9 | ± 0.5 | a | 12.0 | ± 1.0 | a | 19.6 | ± 2.5 | a | 91.1 | ± 9.0 | a |
| ***irx1*** | 1.5 | ± 0.0 | b | 12.7 | ± 0.4 | de | 21.3 | ± 2.8 | b | 23.6 | ± 1.5 | b | 21.9 | ± 2.7 | a | 176.0 | ± 5.2 | b |
| ***irx1 max4*** | 1.5 | ± 0.0 | ab | 11.8 | ± 0.5 | de | 14.6 | ± 1.1 | c | 20.8 | ± 0.7 | c | 18.3 | ± 1.6 | a | 171.8 | ± 14.7 | b |
| ***irx9*** | 1.6 | ± 0.1 | b | 13.3 | ± 0.4 | e | 14.3 | ± 1.0 | c | 23.6 | ± 1.1 | b | 27.7 | ± 2.0 | b | 55.2 | ± 4.4 | c |
| ***irx9 max4*** | 1.8 | ± 0.1 | b | 13.3 | ± 0.4 | e | 15.5 | ± 1.3 | c | 24.6 | ± 0.9 | b | 32.6 | ± 3.6 | b | 55.7 | ± 1.4 | c |

| | Man | | | GalA | | | GlcA | | | Cellulose | | | Acetate | | |
|---|---|---|---|---|---|---|---|---|---|---|---|---|---|---|---|
| | AVG | ± SD | S | AVG | ± SD | S | AVG | ± SD | S | AVG | ± SD | S | AVG | ± SD | S |
| **Col-0** | 9.1 | ± 1.0 | ab | 39.7 | ± 1.9 | ab | 1.9 | ± 1.00 | ad | 357.9 | ± 15.3 | ab | 38.4 | ± 1.5 | a |
| ***tbl29*** | 9.0 | ± 0.7 | ab | 45.4 | ± 4.5 | bc | 3.9 | ± 0.21 | abc | 310.1 | ± 23.4 | bd | 21.5 | ± 0.9 | b |
| ***max4*** | 9.6 | ± 0.7 | ac | 37.9 | ± 2.9 | ab | 3.5 | ± 0.42 | abc | 360.1 | ± 12.3 | a | 39.9 | ± 0.9 | a |
| ***tbl29 max4*** | 8.6 | ± 0.8 | ac | 36.5 | ± 3.7 | ab | 5.0 | ± 0.74 | bc | 362.2 | ± 8.9 | a | 20.3 | ± 1.9 | b |
| ***irx1*** | 12.2 | ± 1.0 | b | 51.2 | ± 2.6 | cd | 5.1 | ± 1.91 | c | 88.0 | ± 5.5 | c | 60.8 | ± 1.7 | c |
| ***irx1 max4*** | 10.4 | ± 0.9 | bc | 49.6 | ± 2.1 | cd | 5.0 | ± 1.10 | c | 118.3 | ± 23.5 | d | 62.1 | ± 2.6 | c |
| ***irx9*** | 21.4 | ± 1.0 | d | 52.8 | ± 2.6 | cd | 1.0 | ± 0.67 | d | 315.7 | ± 15.0 | b | 34.8 | ± 1.9 | d |
| ***irx9 max4*** | 23.0 | ± 1.1 | d | 52.0 | ± 2.2 | d | 0.8 | 0.50 | d | 359.8 | ± 26.4 | a | 34.7 | ± 1.6 | d |

**Supp. Table 2. Effect of *max4* mutation in the cell wall composition of *irx1* and *irx9*.**
Monosaccharide composition, cellulose and acetate content of stem cell walls. Data are represented as mean (AVG) ± the standard deviation (SD) of biological replicates (n=5). Means with different letters are significantly different (Tukey's HSD, $p<0.05$) in the significance (S) column.
Fuc=Fucose; Rha=Rhamnose; Ara=Arabinose; Gal=Galactose; Glc=Glucose; Xyl=Xylose; Man=Mannose; GalA; Galacturonic Acid; GlcA= Glucuronic Acid
